# Development and Effectiveness Testing of a Mobile Health Education Package for Stroke Prevention Among Stroke Survivors

**DOI:** 10.1101/2022.06.12.495788

**Authors:** Marufat Oluyemisi Odetunde, Sekinat Omowumi Ibrahim, Gideon Tolu Oluwagbade, Adesola Christiana Odole, Chidozie Emmanuel Mbada, Morenikeji A. Komolafe

## Abstract

**Background:** Interventions based on social cognitive theory are more patient-centered and focused on principles that can enlighten and encourage adoption of healthy habits. Addressing limited stroke literacy among stroke survivors based on social cognitive theory is an effective means to reduce secondary stroke risk. The use of trending Mobile Health (m-health) devices can be a valuable interventional approach to achieve this through improved stroke literacy.

**Objectives:** The objectives of this study were to develop and test effectiveness of m-health based educational package for stroke prevention among stroke survivors.

**Method:** This was a multimodal methodology involving content development and effectiveness testing using Delphi protocol and pretest-posttest design respectively. Development involved items selection, rating and retention; script writing, translation and recording into an audio and video educational packages. Effectiveness testing involved 30 consenting, consecutively assigned SSVs in each of audio (AIG) and video (VIG) intervention group. Stroke literacy was assessed at baseline, 2nd and 4th week post-intervention. Participants’ socio-demographic data were also collected. Data was summarized using descriptive and inferential statistics. Alpha value was set at p<0.05.

**Results:** Majority of the participants were male (63.3%), over 60 years old (51.7%), hypertensive (83.3%) and had tertiary education (31.7%). Knowledge of stroke risk factors improved between AIG and VIG from baseline (11 23±4 01 and 10.07±3.24) to 2nd week (17 73±0.78 and 15.30±1.78) and 4th week (17.97±0.18 and 16.77±1.01) post-intervention respectively. There were significant differences between the two groups (p<0.01).

**Conclusion:** Mobile-health education based on social cognitive theory effectively improves stroke literacy among SSVs and should be tested among larger samples in the community.

## INTRODUCTION

Stroke constitutes a major challenge to physicians and rehabilitation experts worldwide, with high incidence, mortality, disability rates, and costs. There were 25.7 million stroke survivors (SSVs), 6.5 million stroke deaths and 10.3 million first-ever strokes worldwide. ^1^ Low- and middle-income countries (LMIC) bear the highest burden of stroke with 4.85 million deaths compared to 1.6 million in high-income countries (HIC) in 2013. ^1^ Similarly, the number of disability-adjusted life-years (DALYs) in LMIC was 91.4 million as opposed 21.5 million in HIC. In HIC, availability of good health services and effective strategies for stroke prevention and care are associated with better outcome compared with LMIC where resources for these strategies are limited. ^1^ SSVs have higher probability for unrecognized vascular risk factors, including hypertension and diabetes, than those with no previous history of stroke. ^2^ Consequently, compared with first-time strokes, recurrent strokes are linked with greater morbidity and mortality. ^3^

The risk of recurrence of up to 80% of vascular events after stroke may be prevented by modifying vascular risk factors through medical and behavioral interventions. ^4^ There is high rate of stroke recurrence and attendant physical and cognitive disabilities experienced by up to 50% of survivors, with serious economic and social consequences. ^5^ In spite of this, SSVs and their family members lack sufficient stroke education even after suffering a stroke. ^6, 7^ Hence, prevention and health promotions action on stroke must take precedence, ^7^ as fear of stroke recurrence is a serious concern for SSVs and their families. ^8^ SSVs are therefore an important population to target for educational interventions for stroke prevention.

Stroke literacy, previously defined as knowledge of stroke symptoms and stroke risk factors, is an important component of reducing the risk of recurrent stroke, although improved stroke literacy alone is not sufficient to decrease secondary stroke risk. ^9, 10^ Prevention is the overall goal in stroke education and 80% of the time, strokes can be prevented by providing proper education of the signs of stroke. ^11^ Consequently, guidelines for stroke management identify secondary prevention of stroke as one of the major goals of stroke care^12^ European Stroke Organization, 2008) while specific guidelines also exist for prevention of recurrent stroke. ^3^ Modifiable risk factors are often not effectively managed after a stroke or transient ischemic attack^1^. Evidence suggests that changes to healthcare services on patient education or behavior alone without changes to the organization of care delivery such as professional role revision and multidisciplinary team collaboration had no clinically significant changes in modifiable risk factors for stroke ^13^. However, changes to the organization of healthcare services were associated with meaningful improvements in blood pressure and body mass index. Interventions based on social cognitive theory are more patient-centered, focus on the demand side of health care and based on principles that can enlighten and encourage adoption of healthy habits.

Advancements in technology now provide new platform and direction to care for patients with chronic conditions such as stroke. Educational technologies such as printed materials, electronic materials, music, television and phone calls have been implemented to promote stroke awareness and prevention among the general population, patients, caregivers and health professionals with variable and inconsistent outcomes ^14^. Cultural influence on educational intervention is also an important component ^15^. Thus, the component, means and health professionals involved in delivering educational intervention is important to achieving a favourable outcome ^13, 16^. Sophisticated technologies such as image-based technologies, sensor-based technologies and virtual reality-based tele-rehabilitation systems are far from the reach of many people living in LMICs like Nigeria. However, the use of smart-phone software provides a novel approach to encouraging home and on-the-go health intervention in the form of mobile health (mHealth) ^18, 19^ Mobile health interventions specifically suited for LMICs have been researched among Nigerian health professionals and patients in recent years with evidence of good uptake, especially in urban cities. ^18–21^ Specifically, video based educational interventions have been shown to be more effective than written materials for education and behavioural changes for other chronic diseases. ^22^ However, this has been sparsely explored for stroke education in this context. For obvious reasons, stroke interventions should not only be effective but practical and sustainable in terms of affordability, availability, accessibility and acceptability, the criteria that audio-visual packages possess. ^23^ Therefore, for SSVs who have access to smartphones, the use of audio-visual educational packages on stroke becomes an important interventional tool which may be effective for improving stroke literacy. This study adopted the theory of changes to the organizational interventions of healthcare services through recommended professional role revision and multidisciplinary team collaboration. ^13^ The multidisciplinary team was involved and professional role revised in which physiotherapists took the lead stroke prevention role in this study, which was aimed at development and effectiveness-testing of audio-visual based m-health (AVmHealth) educational package for stroke prevention among SSVs.

## MATERIALS AND METHODS

This was a multimodal methodology involving two stages of content development and effectiveness testing using Delphi protocol and pretest-posttest design respectively.

### Stage 1: Development of an educational package using Delphi Protocol

Desk review of available methods and guidelines in previous studies on stroke education in literature were studied and items were selected from relevant ones ^24, 25^ based on what is applicable to Nigerian context. Thirteen domains and 78 topics were selected to develop the content of the educational package. The domains are medical knowledge, risk factors, treatment with medication, treatment with surgery, treatment with herbal or alternative medicine, rehabilitation, promotion of healthy lifestyle, dietary habits, prevention strategy, special problems, coping, other topics and importance of outpatient clinic follow-up. Five experts in stroke rehabilitation were thereafter invited to partake in the content validity survey. They included two neurologists, two physiotherapists, and one occupational therapist. The recommended number of experts to review an instrument varies from 2 to 20 individuals. ^26^ At least 5 people are suggested to review the instrument to have sufficient control over chance agreement. ^26^

A cover letter was included with the content validity survey, explaining why experts were invited to participate, along with clear and concise instructions on how to rate each item. Each item was rated for selection or exclusion from the educational package by rating based on degree of agreement with relevance of each item. A 4-point Likert scale was used and responses included: 1 = *not relevant*, 2 = *somewhat relevant*, 3 = *quite relevant* and 4 = *very relevant*. Ratings of 1 and 2 are considered content invalid while ratings of 3 and 4 are considered content valid ^26^. Very relevant items were chosen for this study. Item Content Validation Index (I-CVI) is computed as the number of experts giving a rating of “*very relevant*” for each item divided by the total number of experts. Values range from 0 to 1 where I-CVI >0.79, the item is relevant, between 0.70 and 0.79 the item needs revisions, and if the value is below 0.70 the item is eliminated. ^26^ After consulting the five experts, the result of the content validity survey was assessed. Items with values 0.8 and above were selected for the contents of this educational package. Out of the 78 topics and 13 domains that were selected initially, 59 items and 11 domains (Table 1) were rated very relevant and were included in the content of the educational package at the development stage, while 19 items were rated not relevant and were discarded (Table 1, Bolden items). Contents of two out of the 13 domains were rated irrelevant, the domains were therefore excluded (Table 1, Bolden items). These domains were: Treatment with surgery and Treatment with Herbal or Alternative medicine.

**Table 1:** Critical appraisal of the checklist of educational package for prevention of recurrent stroke.

| Domain 1: Medical Knowledge |  |  |  |  |  |
| --- | --- | --- | --- | --- | --- |
|  | How relevant is this item? |  |  |  | How representative is this item? |
| **1. Parts of the brain** |  |  |  |  |  |
| **2. Right and left hemisphere differences** |  |  |  |  |  |
| **3. Location of blood vessels** |  |  |  |  |  |
| 4. Definition of stroke |  |  |  |  |  |
| 5. Types of stroke |  |  |  |  |  |
| 6. Difference between stroke and TIA |  |  |  |  |  |
| 7. Diagnosis of stroke |  |  |  |  |  |
| 8. Signs/symptoms of stroke |  |  |  |  |  |
| 9. Effect of family history |  |  |  |  |  |
| 10. Emergency treatment |  |  |  |  |  |
| 11. Effects of stroke |  |  |  |  |  |
| 12. Complications |  |  |  |  |  |
| 13. Co-morbidities |  |  |  |  |  |
| 14. Recurrence |  |  |  |  |  |

| Domain 2: Risk Factors |  |  |  |  |  |
| --- | --- | --- | --- | --- | --- |
|  | How relevant is this item? |  |  |  | How representative is this item? |
| 15. High blood pressure |  |  |  |  |  |
| 16. Heart disease |  |  |  |  |  |
| 17. Diabetes |  |  |  |  |  |
| 18. Smoking |  |  |  |  |  |
| 19. Alcohol |  |  |  |  |  |
| 20. Cholesterol |  |  |  |  |  |
| 21. Obesity |  |  |  |  |  |
| 22. Exercise |  |  |  |  |  |
| 23. Stress |  |  |  |  |  |

| Domain 3: Treatment With Medication |  |  |  |  |  |
| --- | --- | --- | --- | --- | --- |
|  | How relevant is this item? |  |  |  | How representative is this item? |
| **24. Blood thinners/antiplatelets** |  |  |  |  |  |
| 25. Side effects |  |  |  |  |  |
| 26. Success of medication |  |  |  |  |  |
| **27. Dose** |  |  |  |  |  |
| 28. Reason for medication |  |  |  |  |  |
| 29. Duration of medication |  |  |  |  |  |

Relevant Scale: 1= Not relevant; 2 = Somewhat relevant; 3 = Quite relevant; 4 = Very relevant Representative Scale: 1 = Not Representative; 2 = Useful but not Representative; 3 =Representative
| **Domain 4: Treatment with Surgery** |  |  |  |  |  |
| --- | --- | --- | --- | --- | --- |
|  | **How relevant is this item?** |  |  |  | **How representative is this item?** |
| **30. Deciding surgery** |  |  |  |  |  |
| **31. Diagnostic method** |  |  |  |  |  |
| **32. Recovery** |  |  |  |  |  |
| **33. Cost** |  |  |  |  |  |
| **34. Complications** |  |  |  |  |  |

| **A. Domain 5: Treatment With Herbal or Alternative Medicine** |  |  |  |  |  |
| --- | --- | --- | --- | --- | --- |
|  | **How relevant is this item?** |  |  |  | **How representative is this item?** |
| **35. Difference between Western and African medicine** |  |  |  |  |  |
| **36. Combine herbal medicine and surgery** |  |  |  |  |  |
| **37. Acupuncture** |  |  |  |  |  |
| **38. Diagnostic method** |  |  |  |  |  |

| **Domain 6: Rehabilitation** |  |  |  |  |  |
| --- | --- | --- | --- | --- | --- |
|  | **How relevant is this item?** |  |  |  | **How representative is this item?** |
| **39. General rehabilitation (Physiotherapy, Occupational therapy and Speech therapy)** |  |  |  |  |  |
| **40. Care for bedsores** |  |  |  |  |  |
| **41. Depression** |  |  |  |  |  |
| **42. Anger management** |  |  |  |  |  |
| **43. Behavioural issues** |  |  |  |  |  |
| **44. Sexual activity** |  |  |  |  |  |

| **Domain 7: Promotion of healthy lifestyle** |  |  |  |  |  |
| --- | --- | --- | --- | --- | --- |
|  | **How relevant is this item?** |  |  |  | **How representative is this item?** |
| **45. Leisure activities** |  |  |  |  |  |
| **46. Physical fitness** |  |  |  |  |  |
| **47. Adaptive activities** |  |  |  |  |  |
| **48. Fatigue** |  |  |  |  |  |
| **49. Rest and sleeping patterns** |  |  |  |  |  |
| **50. Engaging in activities of daily living** |  |  |  |  |  |

Relevant Scale: 1= Not relevant; 2 = Somewhat relevant; 3 = Quite relevant; 4 = Very relevant Representative Scale: 1 = Not Representative; 2 = Useful but not Representative; 3 =Representative
| Domain 8: Dietary Habits |  |  |  |  |  |
| --- | --- | --- | --- | --- | --- |
|  | How relevant is this item? |  |  |  | How representative is this item? |
| 51. Meat intake |  |  |  |  |  |
| 52. Fatty food intake |  |  |  |  |  |
| 53. Fish intake |  |  |  |  |  |
| 54. Vegetable and fruit intake |  |  |  |  |  |
| 55. Bean intake |  |  |  |  |  |
| **56. Milk intake** |  |  |  |  |  |
| 57. Salt intake |  |  |  |  |  |

| Domain 9: Prevention strategy |  |  |  |  |  |
| --- | --- | --- | --- | --- | --- |
|  | How relevant is this item? |  |  |  | How representative is this item? |
| 58. Reduce risk |  |  |  |  |  |
| 59. Recognize stroke signs |  |  |  |  |  |
| 60. Respond by calling a doctor |  |  |  |  |  |
| 61. Respond by going to the hospital |  |  |  |  |  |

| B. Domain 10: Special problems |  |  |  |  |  |
| --- | --- | --- | --- | --- | --- |
|  | How relevant is this item? |  |  |  | How representative is this item? |
| 62. Cognition |  |  |  |  |  |
| 63. Aphasia |  |  |  |  |  |
| 64. Bladder/bowel function |  |  |  |  |  |
| 65. Dysphagia |  |  |  |  |  |
| 66. Facial paralysis |  |  |  |  |  |

| Domain 11: Coping |  |  |  |  |  |
| --- | --- | --- | --- | --- | --- |
|  | How relevant is this item? |  |  |  | How representative is this item? |
| 67. Problem solving |  |  |  |  |  |
| 68. Relaxation exercises |  |  |  |  |  |
| 69. Support system |  |  |  |  |  |
| 70. Time management |  |  |  |  |  |
| 71. Coping strategies |  |  |  |  |  |

Relevant Scale: 1= Not relevant; 2 = Somewhat relevant; 3 = Quite relevant; 4 = Very relevant Representative Scale: 1 = Not Representative; 2 = Useful but not Representative; 3 =Representative
| Domain 12: Other Topics |  |  |  |  |  |
| --- | --- | --- | --- | --- | --- |
|  | How relevant is this item? |  |  |  | How representative is this item? |
| 72. Ability to use extremities |  |  |  |  |  |
| 73. Ability to work |  |  |  |  |  |
| **74. Ability to drive** |  |  |  |  |  |
| **75. What people think** |  |  |  |  |  |
| **76. How you will look** |  |  |  |  |  |

| C. Domain 13: Importance of outpatient clinic follow-up |  |  |  |  |  |
| --- | --- | --- | --- | --- | --- |
|  | How relevant is this item? |  |  |  | How representative is this item? |
| 77. Check medical condition |  |  |  |  |  |
| 78. Check ability to use skills learned in rehabilitation |  |  |  |  |  |

The selected items of the educational package were written into a script in English language and translated to Yoruba, the indigenous language of the southwestern Nigeria, the target population for this study. The translation was done by a Yoruba language teacher at a Secondary school level and was validated by a stroke rehabilitation expert who is proficient in both Yoruba and English languages. After the development of the script, it was then made into a speech by an orator and recorded by an audio technician into a 14-minute audio package in English and Yoruba languages. The educational audios were then merged with pictures relevant to the content of the educational package to produce an audio-visual package made available in video MP4 format, playable on mobile phones, laptops, televisions and other MP4 compatible devices.

### Stage 2: Pilot testing of the educational package

Pilot testing of the AVmHealth educational package was conducted over a period of 4 weeks in three phases viz: the pre-intervention phase, the initial post-intervention phase and the final post-intervention phase.

#### Participants

Participants in this study were stroke (SSVs) who were purposively recruited from the Physiotherapy and Neurology outpatient clinics of Obafemi Awolowo University Teaching Hospital (OAUTH), Ile-Ife, Uniosun Teaching Hospital (UTH), Osogbo, and State Hospital Asubiaro (SHA), Osogbo all in Osun state, Nigeria. Stroke survivors with no cognitive impairment (scores of 2 or less on the Short Portable Mental Status questionnaire (SPMSQ)), who can read and/or understand stroke literacy questionnaire and SPMSQ were included in the study; while SSVs with scores of > 2 on the SPMSQ, those on hospital admission or those with active psychiatric disease or dementia were excluded from the study. Participants were divided into two groups viz: audio intervention group (AIG) and video intervention group (VIG).

#### Sample size calculation

The sample size was determined using a general rule of thumb for the large enough sample condition which is n≥30, where n is the sample size. This is based on central limit theorem which states that if a sample size is large enough, the distribution will be approximately normal. The general rule of n≥30 occurs. ^27^ The recommended number of 30 stroke survivors was recruited for each of AIG and VIG group.

### Instrument

i. Short Portable Mental Status Questionnaire (SPMSQ): This is a 10-item questionnaire used to assess the participants’ level of cognitive impairment. Interpretation of the scores is based on the number of errors, with 0-2 errors having normal mental functioning, 3-4 errors having mild cognitive impairment, 5-7 errors having moderate cognitive impairment and 8 or more errors having severe cognitive impairment. ^28^ It has been found reliable and valid in distinguishing individuals with dementia from cognitively intact individuals with sensitivity and specificity of 0.74 and 0.79 for the telephone test and 0.74 and 0.91 for the face-to-face test, respectively. Both test versions correlated significantly with the Mini-Mental State Examination^28^
ii. Stroke literacy questionnaire for stroke survivors (Adapted from Obembe *et al*., ^29^ and Denny *et al*., ^30^). It is a four-section questionnaire that contained socio-demographic and clinical data; general stroke knowledge questions; signs, symptoms and risk factors of stroke and actions to be taken in event of stroke.
iii. Audio-visual mobile health educational package containing evidence based stroke education.

### Procedure

Ethical approval was obtained from the Ethics and Research Committee of Obafemi Awolowo Teaching Hospitals, Ile-Ife, with number, ERC/2021/06/45. Permission to conduct the study was also obtained from the head of department, Medical Rehabilitation, Obafemi Awolowo University Ile-Ife, and the heads of the each clinic where the participants were recruited. A verbal explanation of the purpose and details of the study were given to the participants. The verbal informed consent of each SSV was obtained followed by screening for cognitive impairment with the SPSMQ. At the Pre-intervention phase, stroke survivors with scores of 2 or less on the SPSMQ were included in the pilot testing. Eligible SSVs were subsequently assessed for baseline stroke literacy using the stroke literacy questionnaire (SLQ). The educational package was shared to each SSV in each of AIG and VIG group via their mobile phones at the various clinics. Participants were instructed to use the AVmHealth educational package for at least three times a week and were further informed about the subsequent re-assessment on stroke literacy that took place 2^nd^ and 4^th^ week post intervention. After the first meeting, the researcher called each SSV on phones twice a week for two weeks to follow up and remind them to use the AVmHealth package. At the initial post-intervention phase SLQ was re-administered on participants after two weeks of intervention at their various clinics or through phone calls. The SSVs were encouraged to continue the use of AVmHealth educational package at least thrice in a week for another two weeks. The final post-intervention phase involved administering SLQ on the SSVs after the 4^th^ week at their various clinics or through phone calls.

### Data analysis

Descriptive statistics was used to characterize demographics and clinical characteristics using mean, standard deviation, frequency and percentages. Repeated measure ANOVA was used to compare stroke literacy within group at baseline, two weeks and four weeks post intervention; while mixed method ANOVA was used for between group comparisons. Alpha value was set at P<0.05. The data analysis was carried out using Statistical Package for Social Sciences (SPSS) software version 22.

## RESULT

### Development of the Educational Package

Following the Delphi guidelines, a process that involved item selection, retention, rejection using consensus from experts in stroke rehabilitation, the contents of the educational for stroke prevention among stroke survivors (SSVs) was successfully developed. Out of the 78 topics and 13 domains that were selected initially, 59 items and 11 domains were retained to produce the educational package (Table 1).

### Effectiveness-testing of the Educational Package

#### Socio-demographic characteristics of participants

A total of 60 SSVs, consecutively and equally assigned into AIG and VIG participated in the effectiveness-testing of this educational package. Majority were male (63.3%), over 60 years of age (51.7%) and had tertiary level education (31.7). The details of socio-demographic and clinical characteristics of the participants are presented in table 2.

**Table 2:** Socio-demographic and Clinical Variables of stroke survivors.

| Variables | Frequency | Percentage (%) |
| --- | --- | --- |
| **Gender** |  |  |
| Male | 38 | 63.3 |
| Female | 22 | 36.7 |
| **Age category (Years)** |  |  |
| 41-50 | 11 | 18.3 |
| 51-60 | 18 | 30.0 |
| Over 60 | 31 | 51.7 |
| **Level of Education** |  |  |
| No /Primary Education | 11 | 18.3 |
| Secondary Education | 15 | 25.0 |
| Post-secondary (OND/NCE) | 12 | 20.0 |
| Tertiary HND/Bachelor's | 19 | 31.7 |
| Post-graduate | 03 | 5.0 |
| **Smoking status** |  |  |
| ***Past Smoker*** |  |  |
| No | 54 | 90 |
| Yes | 06 | 10 |
| ***Current Smoker*** |  |  |
| No | 60 | 100 |
| Yes | 0 | 0 |
| **Hypertensive** |  |  |
| No | 10 | 16.7 |
| Yes | 50 | 83.3 |
| **Prior Stroke** |  |  |
| No | 08 | 13.3 |
| Yes | 52 | 86.7 |

#### Comparison of within group differences of participants

Results of within group stroke literacy are presented in tables 4 (VIG) and 5 (AIG). The VIG participants’ stroke literacy at baseline, 2 weeks and 4 weeks post intervention by age group are presented in table 3. There was a general increase in mean scores of participants with age groups <40 having the highest mean score in stroke knowledge at 2^nd^ and 4^th^ week and age group 41-50 having the highest mean score in knowledge of risk factors and symptoms at 2^nd^ week and 4^th^ week. All age group recorded almost perfect scores in the knowledge of actions to be taken at baseline, 2^nd^ and 4^th^ week post-intervention. There was no significant increase in stroke knowledge among the SSVs between age categories. Table 4 shows the results of within group comparison of AIG participants’ scores on stroke literacy at baseline, 2 weeks and 4 weeks by level of education. Participants with post graduate education have the highest mean score at baseline, 2 weeks and 4 weeks post intervention. There is no significant difference in stroke literacy among the SSVs except in stroke knowledge at 4^th^ week post intervention (F= 2.824 p= 0.038). Post hoc analysis further revealed significance difference between primary and secondary levels of education and also between the primary and the post-secondary levels of education.

**Table 3:** Comparison of within group scores for VIG participants by age group.

| Variables | ≤40 | 41-50 | 51-60 | >60 | F-Value | P- Value |
| --- | --- | --- | --- | --- | --- | --- |
|  | **X±SD (n=2)** | **X ±SD(n=4)** | **X ±SD(n=9)** | **X ±SD(n=15)** |  |  |
| **Stroke Knowledge** |  |  |  |  |  |  |
| Baseline | 2.50 ±0.707 | 1.00±0.817 | 1.11±0.782 | 1.40±0.737 | 2.127 | 0.121 |
| 2 weeks | 4.00±0.000 | 2.75±0.500 | 2.89±0.782 | 2.93±0.594 | 1.963 | 0.144 |
| 4 weeks | 4.00±0.000 | 3.50±0.577 | 3.22±0.667 | 2.97±0.594 | 1.076 | 0.377 |
| **Signs, Symptoms and Risk factors of stroke** |  |  |  |  |  |  |
| Baseline | 8.50±0.707 | 10.50±1.732 | 10.56±4.391 | 9.87±3.067 | 0.247 | 0.863 |
| 2 weeks | 15.50±0.707 | 15.75±2.217 | 15.44±2.128 | 15.07±1.668 | 0.182 | 0.908 |
| 4 weeks | 16.50±0.707 | 17.00±1.414 | 16.78±0.972 | 16.73±1.033 | 0.113 | 0.952 |
| **Actions taken** |  |  |  |  |  |  |
| Baseline | 7.00±0.000 | 6.75±0.500 | 6.89±0.333 | 6.93±0.258 | 0.431 | 0.733 |
| 2 weeks later | 7.00±0.000 | 6.75±0.500 | 6.89±0.333 | 6.93±0.258 | 0.431 | 0.733 |
| 4 weeks later | 7.00±0.000 | 6.75±0.500 | 6.88±0.354 | 6.90±0.316 | 0.210 | 0.888 |

**Table 4:** Comparison of within group scores for Audio Intervention Group at baseline, 2 weeks and 4 weeks by level of education.

| **Variables** | Primary Education | Secondary Education | Post-Secondary | Tertiary Education | F-Value | P- Value |
| --- | --- | --- | --- | --- | --- | --- |
|  | X±SD(n=5) | X±SD(n=10) | X±SD(n=6) | X±SD(n=9) |  |  |
| **Stroke Knowledge** |  |  |  |  |  |  |
| Baseline | 1.50±1.00 | 1.30±0.484 | 1.50±0.837 | 1.33±1.000 | 0.778 | 0.550 |
| 2 weeks later | 2.75±0.50 | 3.20±0.632 | 3.00±0.894 | 2.89±0.601 | 0.961 | 0.446 |
| 4 weeks later | 3.25±0.50 | 3.50±0.527 | 3.17±0.753 | 3.44±0.527 | 1.777 | 0.165 |
| **Warning signs and risk factors** |  |  |  |  |  |  |
| Baseline | 8.75±1.708 | 8.80±3.615 | 11.0±2.280 | 10.56±2.698 | 2.769 | 0.049 |
| 2 weeks | 13.75±0.50 | 14.70±1.829 | 15.00±1.789 | 16.56±1.130 | 3.858 | 0.014* |
| 4 weeks | 16.50±1.29 | 16.30±0.949 | 16.83±1.169 | 17.22±0.667 | 1.564 | 0.215 |
| **Actions taken** |  |  |  |  |  |  |
| Baseline | 7.00±00.00 | 6.90±3.160 | 7.00±00.00 | 7.00±00.00 | 0.372 | 0.774 |
| 2 weeks | 7.00±00.00 | 6.90±3.160 | 7.00±00.00 | 7.00±00.00 | 0.372 | 0.774 |
| 4 weeks | 7.00±00.00 | 6.90±3.160 | 7.00±00.00 | 7.00±00.00 | 0.372 | 0.774 |

#### Comparison of between group differences of participants

There are equal numbers of participants (30) in the audio and VIGs with equal gender distribution (19 males and 11 females). The AIG and VIG are not significantly different in terms of socio-demographic variables (Table 5). Similarly, there are no significant associations between the AIG or VIG and the gender, age or level of education (p>0.05). Table 6 presents comparison of scores of AIG and VIG on stroke literacy at baseline, 2 weeks and 4 weeks post intervention. There is a steady increase in scores at all levels between the two groups, with significant difference between the AIG and VIGs on knowledge of stroke risk factors and warning signs (p<0.01).

**Table 5:**
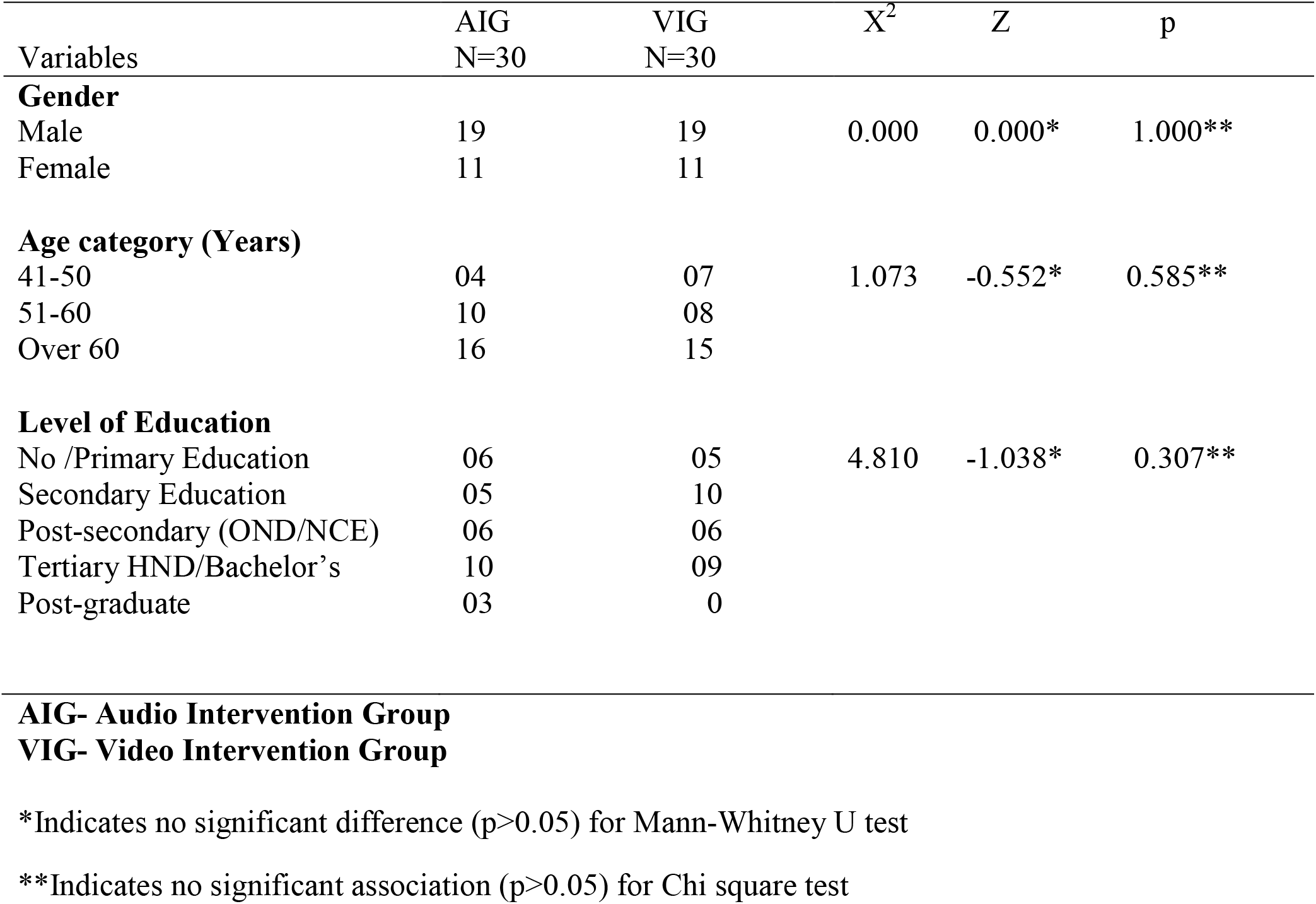
Comparison between Socio-demographic Variables of Audio Intervention Group and Video Intervention Group.

**Table 6:** Comparison of Audio Intervention Group and Video Intervention Group scores on stroke literacy Questionnaire at baseline, 2 weeks and 4 weeks post intervention.

|  | AIG<br>N=30 |  |  | VIG<br>N=30 |  |  |  |  |  |
| --- | --- | --- | --- | --- | --- | --- | --- | --- | --- |
| **Variables** | Baseline<br>X±SD | 2-weeks<br>X±SD | 4-weeks<br>X±SD | Baseline<br>X±SD | 2-weeks<br>X±SD | 4-weeks<br>X±SD | F-value | P-value | Partial<br>Eta<sup>2</sup> |
| Stroke Knowledge | 1.43 (0.94) | 3.33 (0.71) | 3.60 (0.50) | 1.33 (0.80) | 2.97 (0.67) | 3.33 (0.61) | 0.771 | 0.593 | 0.05 |
| Stroke risk factors and<br>warning signs | 11.23 (4.01) | 17.73 (0.78) | 17.97 (0.18) | 10.07 (3.24) | 15.30 (1.78) | 16.77 (1.01) | 12.652 | 0.001 | 0.22 |
| Action taken | 7.00 (0.00) | 7.00 (0.00) | 7.00 (0.00) | 6.90 (0.31) | 6.90 (0.31) | 6.90 (0.31) | - | - | - |
| **AIG- Audio Intervention Group**<br>**VIG- Video Intervention Group** |  |  |  |  |  |  |  |  |  |

#### Comparison of mean scores of AIG and VIGs

Figure 1 shows a steady increase in knowledge of stroke between both AIG and VIGs. Knowledge of stroke risk factors and warning signs for the AIG and VIGs are represented in figure 2. The mean score of the AIG is higher with the highest difference in the scores at the 2^nd^ week. Action to be taken in the event of stroke by each of AIG and VIG is represented in figure 3. The mean score is constant from baseline, 2^nd^ week and 4^th^ week for the two groups.

**Figure 1:**
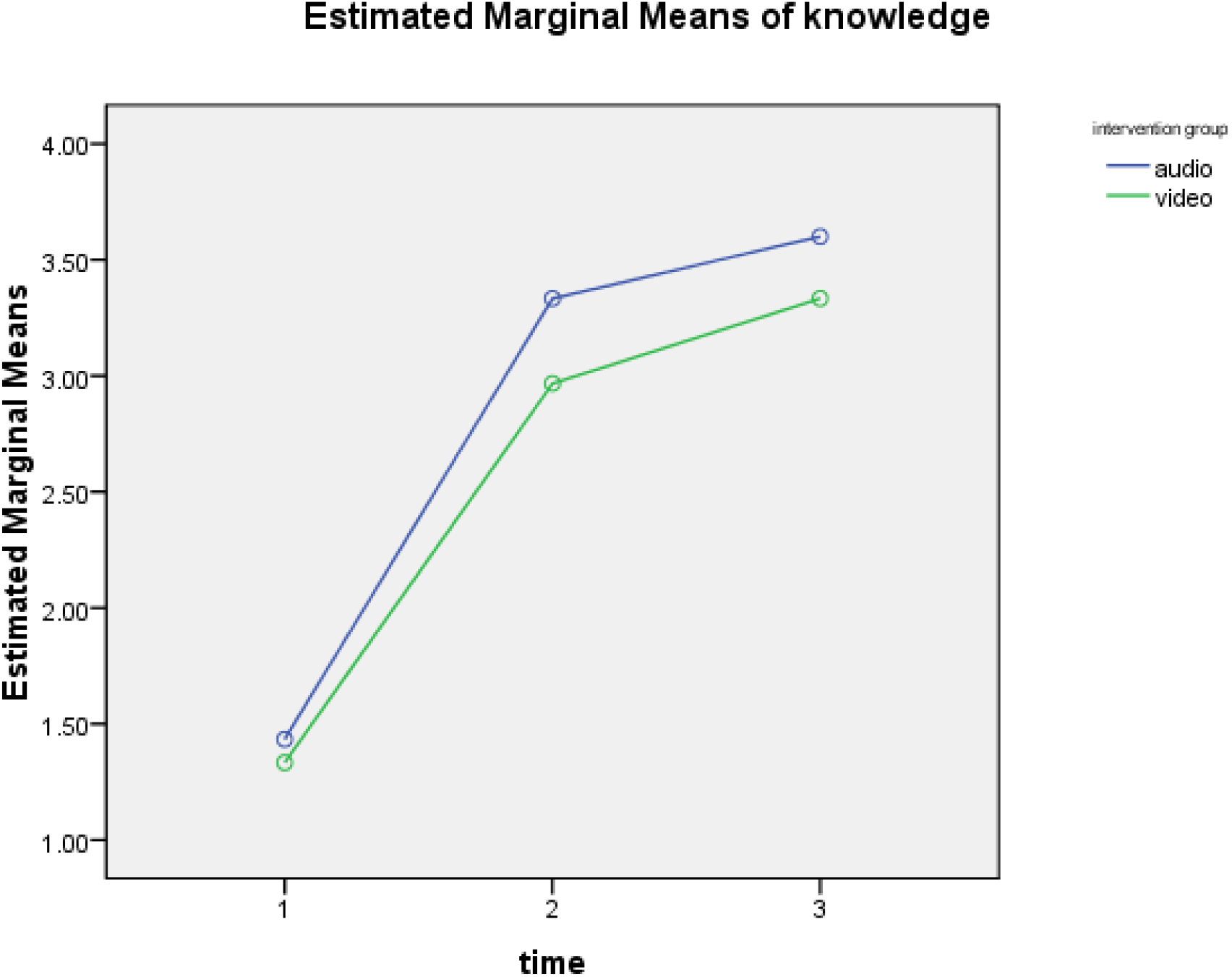
Mean scores on knowledge of stroke between audio and video group at baseline, 2^nd^ and 4^th^ week post intervention

**Figure 2:**
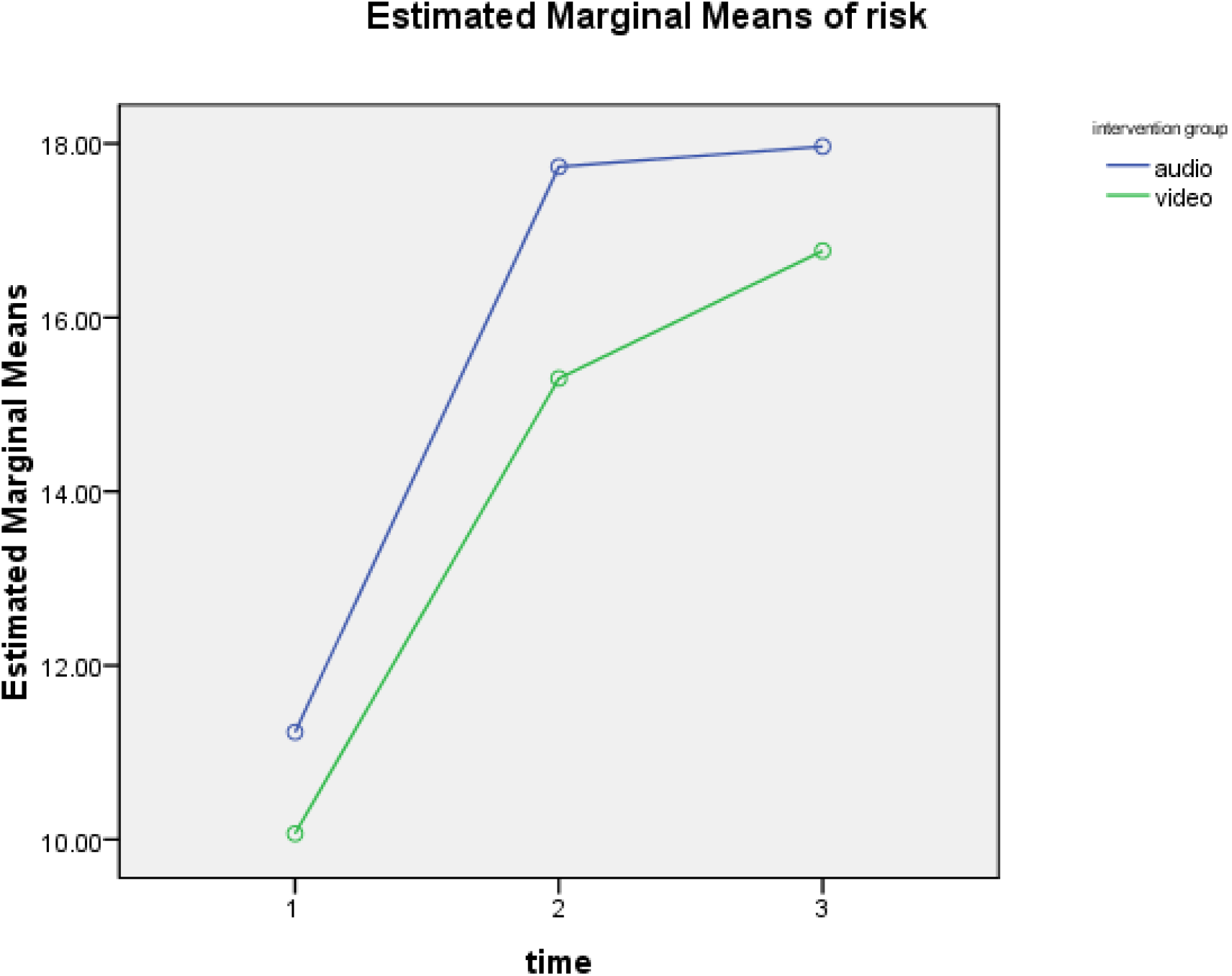
Mean scores of stroke risk factors and warning signs between audio and video group at baseline, 2^nd^ and 4^th^ week post intervention

**Figure 3.**
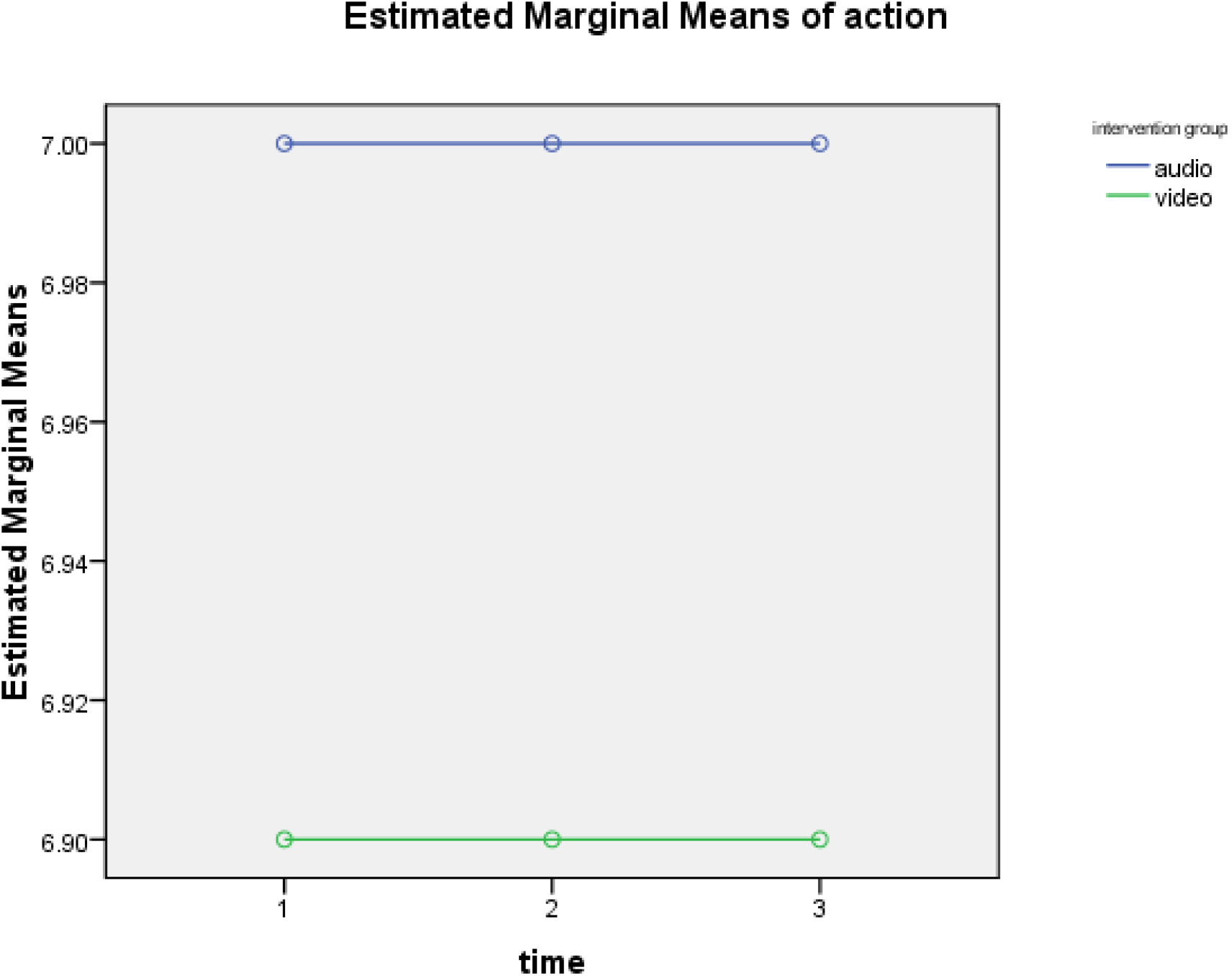
Mean scores on action to be taken between audio and video group at baseline, 2^nd^ and 4^th^ week post intervention

## DISCUSSION

Health professionals need to be aware of the importance of meeting the educational needs of stroke survivors and provide information about all aspects of stroke. SSVs and their caregivers need to be educated via various means of information dissemination such as videos, handbooks, and modern technology. ^24^ In this study, an audio-visual based mobile health (AVmHealth) educational package was developed for stroke prevention among stroke survivors. Knowledge of stroke risk factors is expected to substantially contribute to secondary stroke prevention by improving stroke literacy among stroke survivors, especially in Nigeria and other LMICs where stroke survivors have been reported to lack knowledge of stroke, its risk factors and warning signs. ^6, 31, 32^

The contents of this educational tool are in line with the themes featured in the study of Maniva et al., ^14^. The tool was validated by stroke rehabilitation experts, consistent with previous studies in which a new tool was developed and the items were assessed for content validation. ^33, 34^ The development of AVmHealth educational package in this study was conceptualized and initiated by a physiotherapist and involved a multidisciplinary team. This corroborates the recommendation of professional role revision and multidisciplinary team collaboration emphasized by Joel et al., ^13^ with documented evidence of meaningful outcome. It was a form of engaging material delivered via mobile phones which is more convenient for participants. The intervention was based on social cognitive theory which is more patient-centered and provided information that enlightened and encouraged stroke survivors to adopt healthy habits. The choice of mobile phones as an easy means of information delivery is supported by similar reports of no difficulty with operating the exercise package on the mobile device or at home. ^32, 35^ The educational tool in this study was of 14-minute duration, in contrast to a stroke literacy video of 5-minute duration by Denny et al.,^9^ and a 30-minute video home-based exercise for SSVs by Odetunde et al., ^36^ The varied duration was to address the study objectives and accommodate the contents of each of the videos. This study revealed that the AVmHealth stroke education was associated with improved stroke literacy at the 2^nd^ and 4^th^ week post intervention, consistent with previous studies on stroke education. ^30, 37^ In the AIG, stroke education were associated with improved stroke knowledge and recognition of stroke risk factors and warning signs at the 2^nd^ and 4^th^ week post intervention. The knowledge improved significantly across age group in stroke risk factors and warning signs as well as across level of education in stroke knowledge. In the VIG however, the increase in stroke knowledge is not statistically significant across age group and between genders. Only the levels of education showed significant increase with identification of stroke risk factors and warning signs, between the primary and the tertiary levels of education 2^nd^ week post-intervention. These findings are consistent with previous studies on stroke education of Denny et al., ^30^ and Maisarah et al., ^36^. Earlier studies, have reported increase in stroke knowledge across education levels, with the largest effect on those with at least college education. ^38, 39^

The results of higher mean scores of the AIG than the VIG in all aspects of stroke literacy assessed and at all points is surprising, as naturally the VIG with added advantage of visual feedback is expected to score higher. This finding is supported by earlier reports of many advantages of information dissemination through media such as radio and mobile phones including easy availability to a large population which offer health promoters many advantages over other means. ^40^ Access to health information is an important way to reduce the social and economic impact of preventable and non-preventable diseases and illnesses. ^41^. Availability and accessibility of audio may explain why the use of audio material is common in this environment in the form of music, spiritual messages and public information via radios and pre-recorded audio packages. The AVmhealth based educational package for this study was developed as a form of an engaging material and was delivered via mobile phones. This is in agreement with submission of Angula et al., ^42^ that mobile devices have become the most dominant tools to disseminate information worldwide nowadays and that a large number of people will benefit whether they attend clinics or not, once they have smart phones. Considering the peculiar situation of LMICs, challenges of poor internet facility and erratic power supply that limits the uptake of mHealth via the use of mobile telephone and other advanced technological devices was bypassed in the audio-visual based m-Health educational package employed in this study. This allows for possibility of uptake of this tool among rural and urban poor population of Nigeria and other LMICs. Another positive implication from this study is that video technology can be used as a substitute for conventional education to improve stroke literacy among SSVs. Finally, video-based educational intervention on stroke is a novel approach in clinical practice in Nigeria. The fact that the positive outcomes obtained in this study corroborate similar studies serves as a stimulus for wider use of video-based technology in the dissemination of educational intervention materials among SSVs.

In conclusion, AVmhealth educational package is useful, understandable, effective in improving stroke literacy among stroke survivors and may be effective for secondary stroke prevention. The tool should be tested among larger sample in different settings in the community.

## Authors’ contribution

M.O conceptualized the study, conducted the research, prepared the manuscript and took part in data analysis and interpretation. S.O and G.T were involved in conceptualization of the study, data collection, analysis and interpretation. A.C was involved in the research design and proofread the manuscript for intellectual contribution. C.E was involved in research design, data analysis and interpretation and proof-reading of the manuscript for intellectual contribution; M.A was involved in research design and proof reading. All authors read and approved the final manuscript.

## Acknowledgements

The authors are thankful to all the stroke survivors who participated in this study. We are also grateful to Dr Olamide Olaoye of the Department of Medical Rehabilitation, Obafemi Awolowo University, Ile-Ife, Nigeria for helping with data analysis.

## Data availability statement

Data generated from this study are available on reasonable request from the corresponding author

